# Heterochrony in *orthodenticle* expression is associated with ommatidial size variation between *Drosophila* species

**DOI:** 10.1101/2021.03.17.435774

**Authors:** Montserrat Torres-Oliva, Elisa Buchberger, Alexandra D. Buffry, Maike Kittelmann, Genoveva Guerrero, Lauren Sumner-Rooney, Pedro Gaspar, Georg C. Bullinger, Javier Figueras Jimenez, Fernando Casares, Saad Arif, Nico Posnien, Maria D. S. Nunes, Alistair P. McGregor, Isabel Almudi

## Abstract

The compound eyes of insects exhibit extensive variation in ommatidia number and size, which affects how they see and underlies adaptations in their vision to different environments and lifestyles. However, very little is known about the genetic and developmental bases of differences in compound eye size. We previously showed that the larger eyes of *Drosophila mauritiana* compared to *D. simulans* is generally caused by differences in ommatidia size rather than number. Furthermore, we identified an X-linked chromosomal region in *D. mauritiana* that results in larger eyes when introgressed into *D. simulans*. Here, we used a combination of fine-scale mapping and gene expression analysis to further investigate positional candidate genes on the X chromosome. We found earlier expression of *orthodenticle (otd)* during ommatidial maturation in third instar larvae in *D. mauritiana* than in *D. simulans*, and we show that this gene is required for the correct organisation and size of ommatidia in *D. melanogaster*. We discovered that the activity of an *otd* eye enhancer is consistent with the difference in the expression of this gene between species, with the *D. mauritiana* enhancer sequence driving earlier expression than that of *D. simulans*. We also identified potential direct targets of Otd that are differentially expressed between *D. mauritiana* and *D. simulans* during ommatidial maturation. Taken together, our results suggest that differential timing of *otd* expression contributes to natural variation in ommatidia size between *D. mauritiana* and *D. simulans,* which provides new insights into the mechanisms underlying the regulation and evolution of compound eye size in insects.

## Introduction

Understanding the genetic basis of phenotypic diversity is one of the central themes of evolutionary developmental biology. While the causative genes and even mutations underlying evolutionary changes in a growing list of phenotypes have been identified (e. g. (Arif et al., 2013b; Arnoult et al., 2013; Klaassen et al., 2018; Ramaekers et al., 2019; Ridgway et al., 2024; Santos et al., 2017; Xia et al., 2024)) and see (Courtier-Orgogozo et al., 2020) for a more comprehensive list), we still know relatively little about the genetic basis for the evolution of organ size.

Insects exhibit remarkable variation in the size and shape of their compound eyes, which has allowed these animals to adapt to different environments and lifestyles (Kittelmann and McGregor, 2024; Land and Nilsson, 2012). This variation greatly affects optical parameters and visual sensation, such as the detection of different intensities, polarization and wavelengths of light to varying degrees of contrast sensitivity and acuity (Land and Nilsson, 2012). Compound eyes vary in the size and/or number of ommatidia: wider ommatidia capture more light, which can increase contrast sensitivity; however, larger interommatidial angles can lead to decreased acuity (Land, 1997; Land and Nilsson, 2012). Conversely, having many small ommatidia with narrow interommatidial angles can enhance acuity, but this may decrease contrast sensitivity (Currea et al., 2018; Palavalli-Nettimi and Theobald, 2020; Warrant and Nilsson, 2006; Warrant, 1999).

Differences in ommatidia number and size, as well as trade-offs between these structural features of compound eyes, have been described for a range of different insects (Buffry et al., 2024; Duncan et al., 2021; Gonzalez-Bellido et al., 2011; Horridge, 1977; Posnien et al., 2012; Wakakuwa et al., 2007). Furthermore, variation in ommatidia size across the eye within species is also widely documented (Gaspar et al., 2020; Gonzalez-Bellido et al., 2011; Land, 1989; Perl and Niven, 2016; Streinzer et al., 2013). This size variation suggests areas of regional specialisation, where different visual tasks rely on different parts of the eye.

Several studies have also found extensive variation in eye size within and between closely related species of *Drosophila,* caused by differences in ommatidia number and/or ommatidia area (Arif et al., 2013a; Buchberger et al., 2021; Buffry et al., 2024; Gaspar et al., 2020; Hilbrant et al., 2014; Keesey et al., 2019; Norry and Gomez, 2017; Posnien et al., 2012; Ramaekers et al., 2019; Reis et al., 2020). Despite the pervasive variation in eye morphology and the detailed knowledge about eye development in *Drosophila* (Casares and Almudi, 2016; Casares and McGregor, 2021; Kittelmann and McGregor, 2024; Kumar, 2018), little is known about the genetic and developmental bases for variation in eye size even among *Drosophila* species with very few exceptions (e.g. (Buchberger et al., 2021; Ramaekers et al., 2019).

We previously showed that *D. mauritiana* has larger eyes than *D. simulans* mainly due to larger ommatidia (Arif et al., 2013a; Posnien et al., 2012). Quantitative trait loci (QTL) mapping of this difference identified a large-effect QTL that explains 33% of the species difference (Arif et al., 2013a). Introgression of this X-linked region from *D. mauritiana* into *D. simulans* increased the eye size and ommatidial size of the latter species (Arif et al., 2013a).

Here, we combine higher resolution mapping of this previously characterised X-linked QTL, with transcriptomic analysis of eye-antennal imaginal discs (EADs) of *D. simulans* and *D. mauritiana,* to identify positional candidate genes that are differentially expressed in the developing ommatidia between these two species. We observed a temporal difference in the onset of expression of one of these candidates, the homeobox gene *orthodenticle (otd)*, and we confirmed that Otd is involved in ommatidia organization and size determination. We then carried out ATAC-seq to compare putative regulatory regions of *otd* that may underlie the difference in eye expression of this gene between *D. mauritiana* and *D. simulans.* Our results indicate that differential activity of an orthologous eye enhancer of *otd* between these species results in earlier expression of this homeobox gene during ommatidial maturation in *D. mauritiana*. We hypothesise that this heterochrony in *otd* expression and consequently longer exposure to this transcription factor (TF) in maturing ommatidia in *D. mauritiana* contributes to the development of larger ommatidia in this species.

## Results

### Enlarged ommatidia in *D. mauritiana*

We previously found that the central ommatidia of *D. mauritiana* eyes have wider diameter than those of *D. simulans* (Arif et al., 2013a; Posnien et al., 2012). To examine whether this phenotypic difference is prevalent in all ommatidia across the eye, we imaged the eyes of a female *D. mauritiana* TAM16 and *D. simulans y, v, f* using synchrotron radiation micro-CT (SRµCT) and measured the facet diameter of ommatidia in different regions of the eye using 3D reconstructions (Fig. 1). We corroborated that while the number of ommatidia is similar between these strains of *D. mauritiana* and *D. simulans* the former has larger facets, as observed for other strains of these two species (Buffry et al., 2024). This trend is consistent across anterior, central and posterior facets but is particularly pronounced in the antero-ventral region of the eye (Fig. 1 and Data S1 (Arif et al., 2013a; Posnien et al., 2012)) again as seen for other strains (Buffry et al., 2024).

**Figure 1.**
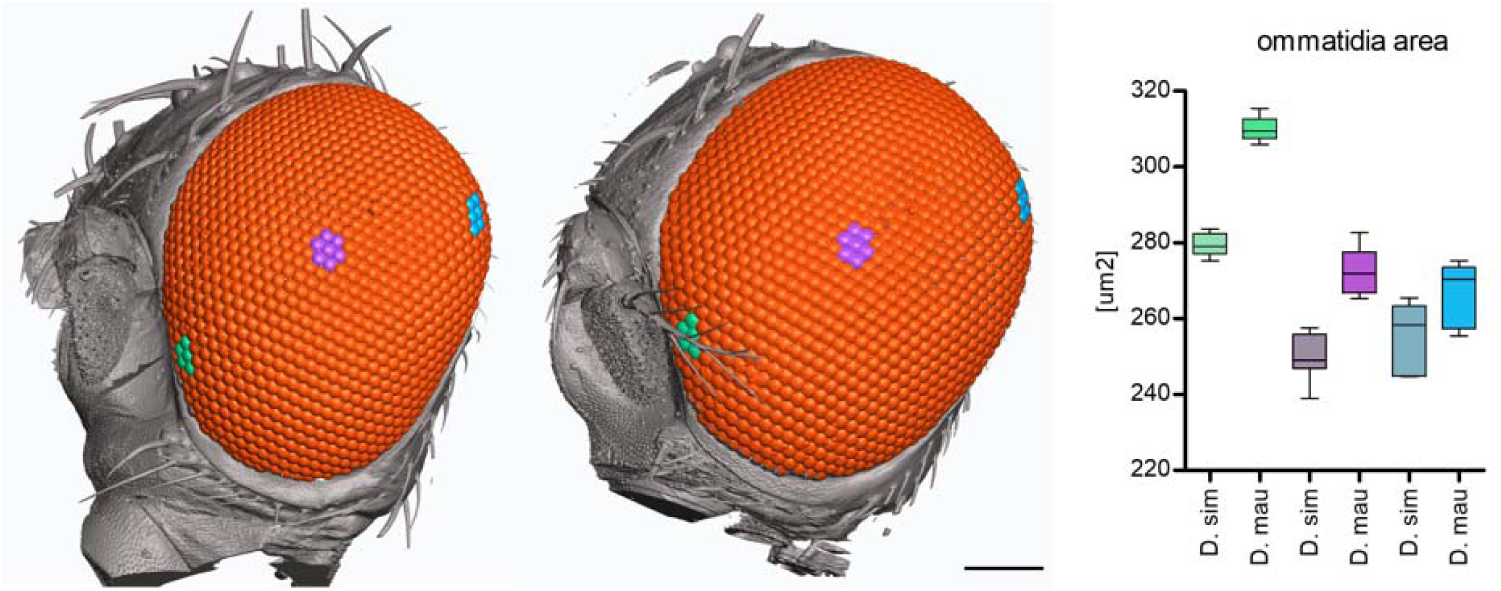
3D reconstruction and ommatidia size measurements from SRµCT data of female *D. simulans* (left) and *D. mauritiana* (right). Facet areas of the ommatidia highlighted in the antero-ventral (green), central (purple) and dorsal-posterior (blue) region of the eye are plotted in corresponding colours (far right). Ommatidia number is 996 for the *D. simulans y, v, f* and 1018 for the *D. mauritiana* TAM16. Scale bar is 100 µm.

### Differentially expressed genes in a candidate region on the X chromosome

Previously we detected a QTL region located between 2.6 Mb and 8 Mb on the X chromosome, which is responsible for 33% of the difference in ommatidia size (Arif et al., 2013a). Furthermore, introgression of approximately 8.3 Mb of this X-linked region (between the *yellow* (*y*) and the *vermillion* (*v*) loci) from *D. mauritiana* TAM16 into *D. simulans y, v, f* significantly increased the eye size of the latter, consistent with the direction of the species difference (Arif et al., 2013a). Further analysis of recombinant males with breakpoints within the introgressed region revealed significant genotype-phenotype associations towards the distal end of the introgressed region near marker *v*, providing a conservative interval of about 2 Mb wherein the X QTL is likely to reside (Fig. 2A). To map the candidate region to higher resolution we generated introgression lines with breakpoints in the 2 Mb interval and compared eye area and central ommatidia diameter of *y*, *f* male progeny (with some *D. mauritiana* DNA in the 2 Mb interval) to that of their *y, v, f* sibling males (*i.e.*, without *D. mauritiana* DNA). We found that *y, f* males had significantly larger eye size than their *y, v, f* siblings in introgression lines IL9.1a (one tailed t=1.80, df=11, p=.026), IL9.1b (one tailed t=1.80, df=11, p<.001) and IL9.2 (one tailed t=1.80, df=11, p=.035) but ommatidia diameter was only significantly different for IL9.1a (one tailed t=1.80, df=11, p=.014) and b (one tailed t=1.80, df=11, p=.005) (Fig. 2A). Ommatidia number and body size did not differ between *y*, *f* males and their respective *y, v, f* sibling males for any of the IL lines (Data S2). These data suggest that the candidate QTL is located in a maximum region of about 662 kb (ChrX: 7,725,195-8,387,618 in *D. simulans*).

**Figure 2.**
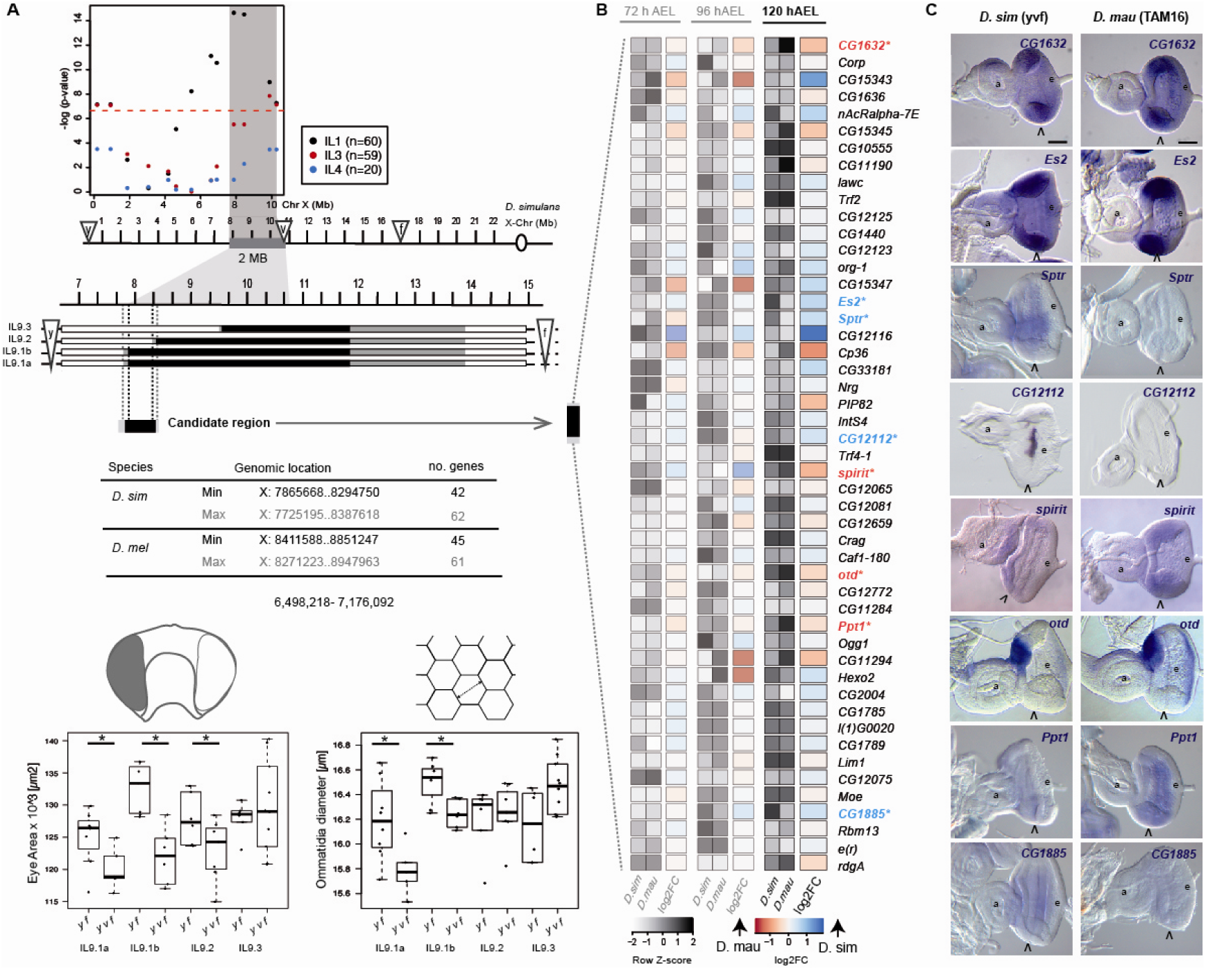
Differential and spatial gene expression. **(A)** Fine-scale mapping of the X chromosome QTL. Marker-phenotype association in male recombinant progeny from three replicate introgression lines (IL1, 3, and 4, single-marker ANOVA analysis). Red dashed line indicates the Bonferroni corrected significance threshold of 0.05. Shaded grey area represents a conservative interval of ∼ 2 Mb encompassing the X linked QTL. Recombination breakpoints of the new introgression lines (IL9.1-9.3) on the X chromosome (shown for *D. simulans* Flybase R2.02) define the 662 kb candidate region. White, black, and grey boxes indicate DNA regions from *D. simulans y, v, f*, *D. mauritiana* TAM16 or not determined, respectively (the latter define the maximum candidate region). The table indicates the number of protein coding genes that are present in the candidate region in *D. simulans* and *D. melanogaster*. Distribution of eye area (left) and ommatidia diameter (right) measurements by genotype and introgression line. Asterisks indicate significance differences between genotypes where *p* <.05 (Data S2). **(B)** Differential expression of 49 protein coding genes located in the introgressed region from (A) and expressed at 72 h AEL, 96 h AEL and 120 h AEL. Genes with significantly higher expression at 120 h AEL in *D. simulans* are highlighted in blue. Genes significantly upregulated in *D. mauritiana* at 120 h AEL are shown in red. **(C)** Expression of differentially expressed genes at 120 h AEL in L3 EADs of *D. simulans* and *D. mauritiana.* Open arrowheads indicate the MF, a: antenna, e: eye in (C).

This mapped region contains 62 protein coding positional candidate genes. To assay which of these candidates are expressed during the generation of ommatidia, we performed RNA-seq experiments on the EADs of 3rd instar larvae (L3). We extracted RNA from *D. mauritiana* and *D. simulans* EADs at three different developmental points: at 72 hours after egg laying (AEL; late L2, at the onset of differentiation marked by the morphogenetic furrow (MF)), when the eye primordium is proliferating and specification of the ommatidial cells has not yet started; at 96 h AEL stage (mid L3) when the MF has moved about half way through the eye disc and the most posterior ommatidia are already determined, and at 120 h AEL (late L3), when most ommatidia are already determined while their final size, structure and shape are being arranged (Torres-Oliva et al., 2018, 2016).

Comparison of the RNA-seq data among these three developmental timepoints showed that transcriptomes of 72 h AEL EADs were the most different in comparison to transcriptomes from both 96 h AEL and 120 h AEL for both species (Fig. S1 and Data S3). This reflects the distinctive processes that are occurring at these developmental stages (Torres-Oliva et al., 2018). We next focused on the expression of genes located within the mapped 0.66 Mb X-linked region at 120 h AEL because at this timepoint ommatidia at different stages of assembly are arranged from posterior to anterior with the most mature, with a full complement of cells, beginning to adopt their final size at the posterior of the eye disc. Of the 62 genes located in this region, 49 were expressed at least at one of the RNA-seq timepoints and only eight of these genes were differentially expressed between these two species at 120 h AEL (Data S3): *spirit, otd* and *Ppt1* showed higher expression in *D. mauritiana*, whereas *CG1632, Es2, Sptr, CG12112* and *CG1885* were more highly expressed in *D. simulans* (Fig. 2B).

We next performed *in situ* hybridization experiments of these eight candidate genes to investigate if they are expressed in the eye field where the ommatidia are being assembled. These assays were carried out in both *D. mauritiana* and *D. simulans*, which allowed us to determine whether the differences in expression levels observed in the RNA-seq datasets are related to differences in spatial expression (Fig. 2C). *Sptr, CG12112* and *spirit* had no detectable expression in the relevant region posterior to the MF (Fig. 2C). *Ppt1* and *CG1885* were expressed both anterior to and immediately posterior to the MF. *CG1632* and *Es2* were ubiquitously expressed in the eye disc, with no clear regional differences. Finally, as previously shown in *D. melanogaster*, *otd* was expressed in the ocellar region of the EAD and in the most posterior region of the eye field (Vandendries et al., 1996). At 120 h AEL *otd* is already expressed in several rows of the most posterior ommatidia of *D. mauritiana* eye discs, whereas *otd* expression is undetectable in the most posterior region of the eye discs of *D. simulans* (Fig. 2C). These results were consistent with our differential expression analysis, as the analyses of spatial expression showed qualitative differences in expression levels for most of the investigated genes. Taken together, these results showed that *otd* is the only differentially expressed positional candidate gene that is expressed in maturing ommatidia (Fig. 2C).

### Differences in *otd* gene expression during eye development between *D. simulans* and *D. mauritiana*

Our results suggested that *otd* transcription in the maturing ommatidia initiates earlier in *D. mauritiana* than in *D. simulans* (Fig. 2C). To investigate this further, we performed additional *in situ* hybridizations at 110 h AEL to compare the onset of *otd* expression in the developing ommatidia of these two species. At this developmental stage, we found that *otd* is already transcribed in *D. mauritiana* eye discs, whereas there was no detectable expression in *D. simulans* discs (Fig. 3A-B). To confirm this heterochrony in *otd* expression we performed immunostainings against Otd protein in developing eye discs (Fig. 3C-D). We measured the number of ommatidial rows that were already specified (i.e. with positive Elav staining) as a proxy of developmental stage and then which of these ommatidial rows showed Otd expression. We observed that *D. mauritiana* eye discs displayed more Otd-positive ommatidia than *D. simulans* eye discs at the same stage (Elav positive ommatidia rows, Fig. 3E, Data S4, *F _1,47_ = 30.3, p-value=1.48 x 10^-06^*). Thus, cells in maturing ommatidia are exposed to the action of Otd for longer in *D. mauritiana* since the expression of this protein extends into the pupal stage of both species (Fig. S2).

**Figure 3.**
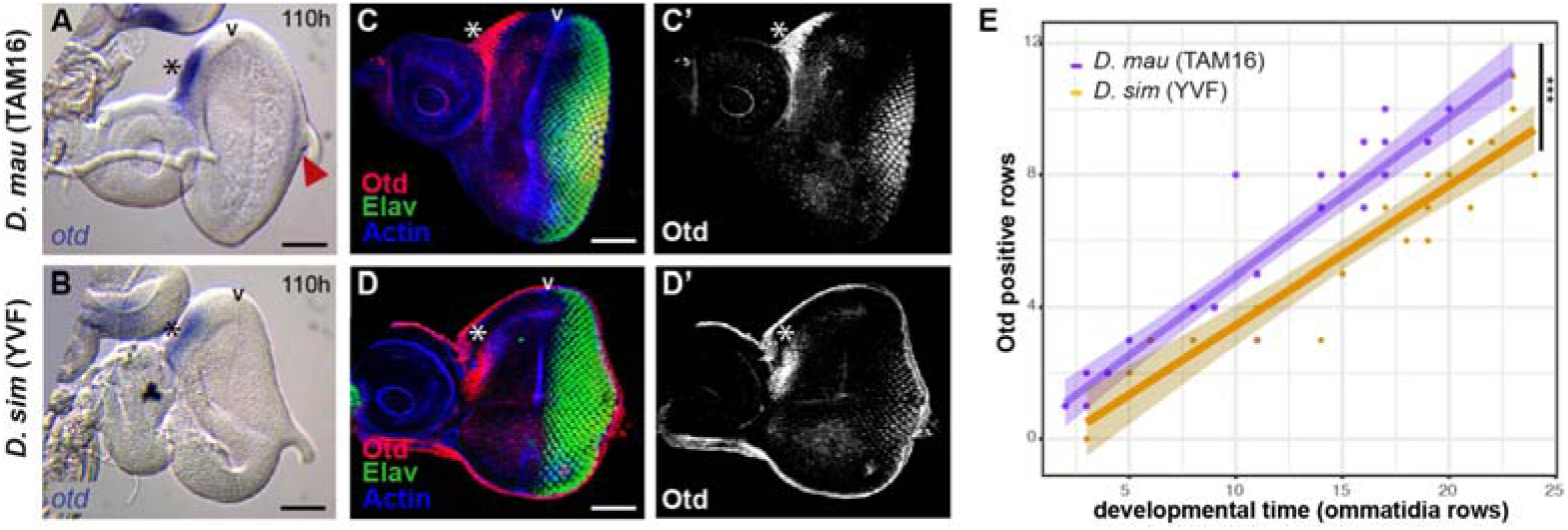
*otd* expression in L3 eye imaginal discs. **(A-D)** *otd* mRNA at 110 h AEL in *D. mauritiana* (A) and *D. simulans* (B). Arrowheads indicate the morphogenetic furrow. Asterisks indicate expression in the ocellar region. *D. mauritiana* already exhibits *otd* mRNA at 110 h (red arrowhead). **(C-D’)** Immunostaining showing Otd protein (magenta, C’, D’) in mature ommatidia (marked in green by Elav) and the ocellar region (asterisk) in *D. mauritiana* (C-C’) and *D. simulans* (D-D’). Staining against actin (blue) was used to mark the morphogenetic furrow. **(E)** Plot showing the number of Otd-positive ommatidia rows (y-axis) at different developmental time points (x-axis, developmental points inferred by number of ommatidia rows). Asterisks represent statistical significance p< 0.001. Scale bars: 50 µm. Anterior regions of the EADs are on the left and posterior on the right.

### *otd* is required for the correct arrangement and size of ommatidia in *Drosophila*

It was previously shown in *D. melanogaster* that *otd* is expressed in photoreceptor cells and required during pupal stages for morphogenesis of their rhabdomeres, and subsequently rhodopsin expression, as well as for the synaptic-column and layer targeting of the photoreceptors (Fichelson et al., 2012; Mencarelli and Pichaud, 2015; Vandendries et al., 1996). We carried out further analysis of *otd* function during eye development using RNAi knockdown and by generating mitotic clones of homozygous *otd* mutant cells. Decreasing *otd* mRNA by overexpressing an *otd-miRNA* construct in cells posterior to the MF (Wang et al., 2010), resulted in defects in the final ommatidial organisation. These defects were rescued by adding a copy of *UAS-otd* (Fig. S3A-C), supporting the specificity of the RNAi-mediated *otd* attenuation. Loss of *otd* in clones resulted in disorganised ommatidia with perturbed shapes and sizes – often smaller than the ommatidia of controls (Fig. S3D). These results show that *otd* expression in the photoreceptor cells of maturing ommatidia is required for the proper regulation of ommatidial organisation and size.

### Differences in chromatin accessibility in the *otd* locus during eye development between *D. simulans* and *D. mauritiana*

Our mapping and expression analyses indicate that the differences in *otd* expression likely contribute to differences in ommatidia size between *D. simulans* and *D. mauritiana.* Given that there is no amino acid difference in the Otd homeodomain between our focal strains of *D. simulans* and *D. mauritiana* (Fig. S4), our data suggest that the causative changes reside in *otd* regulatory regions. Due to the microsyntenic conservation between *D. melanogaster, D. simulans* and *D. mauritiana,* we considered the regulatory landscape of *otd* as the region between its two flanking genes, *Caf1-180* and *CG12772*, revealed by the presence of a Topological Associated Domain (TAD) in the corresponding *D. melanogaster* region (a region of 69 kb in *D. mauritiana* and 70 kb *D. simulans,* Fig. 4A, Fig. S5; http://chorogenome.ie-freiburg.mpg.de/).

**Figure 4.**
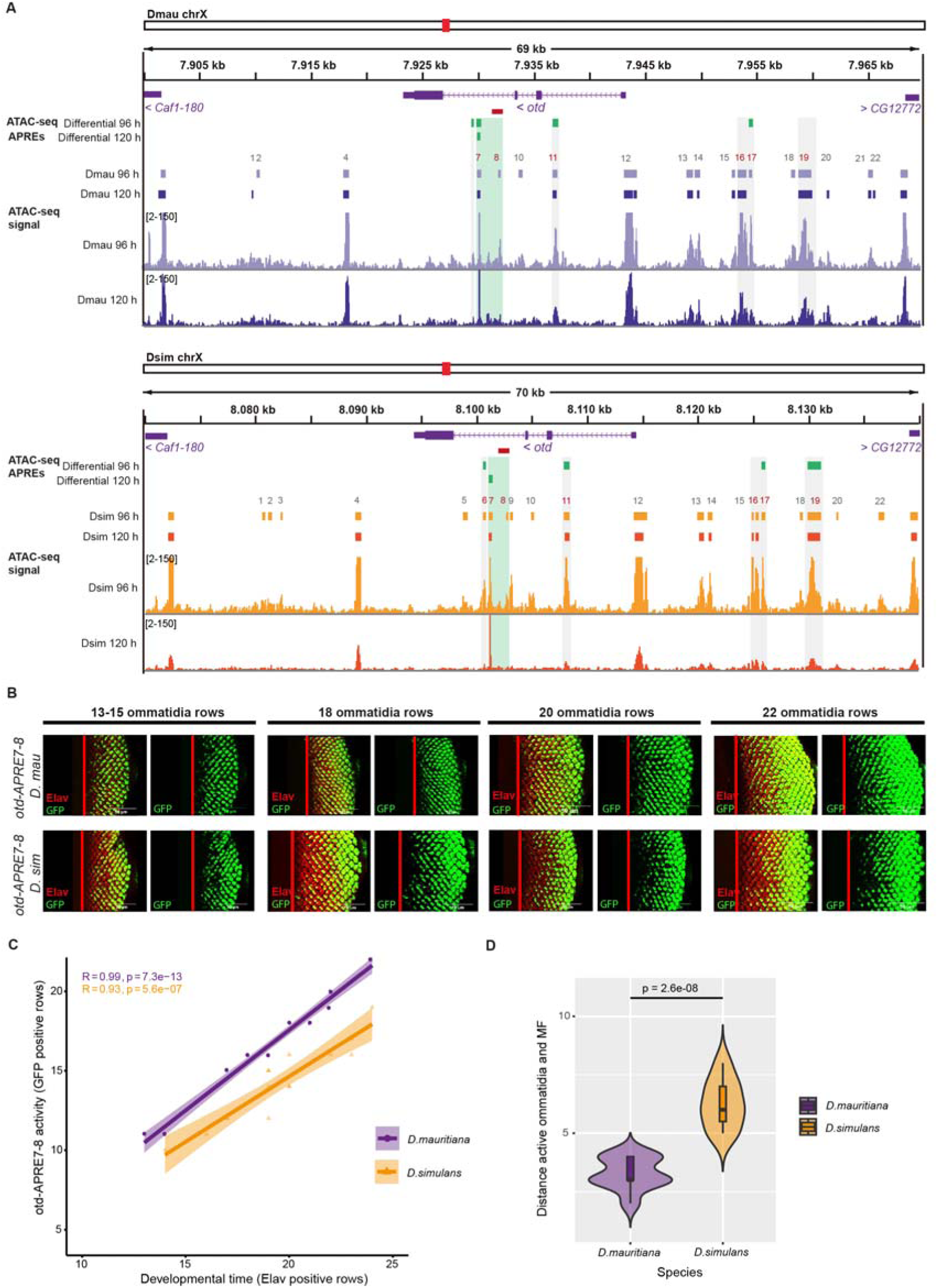
Chromatin accessibility at the *otd* locus. **(A)** Chromatin accessibility profiles at the *otd* locus at 96 h AEL and 120 h AEL in *D. mauritiana* and *D. simulans* EADs. Purple and orange boxes indicate APREs detected in *D. mauritiana* and *D. simulans,* respectively, when mapped against their own reference genome. Green boxes highlight APREs with differential accessibility, identified by re-mapping and peak calling of *D. mauritiana* datasets against *D. simulans* reference genome and *D. simulans* datasets against *D. mauritiana* reference genome, through the "liftOver" tool and MACS2. Red boxes represent the region corresponding to the *D. melanogaster otd^uvi^* allele region. **(B)** *D. simulans* and *D. mauritiana otd-APRE7-8* enhancer activity in 3^rd^ instar larvae eye imaginal discs. **(C)** Plot showing rows with full enhancer activity (y-axis) activated by *otd-APRE7-8* at different developmental time points (x-axis, developmental points inferred by number of ommatidia rows) for both species. **(D)** Violin plots showing the distance between the morphogenetic furrow and the rows with full *otd-APRE7-8* activity (first row counting from the posterior margin of the disc with more than 90% of the ommatidia positive for GFP signal). *D. mauritiana otd-APRE7-8* is active earlier during the differentiation of ommatidia.

To investigate the regulation of *otd* in the developing eyes of *D. simulans* and *D. mauritiana* further, we performed ATAC-seq (Buchberger et al., 2021; Buenrostro et al., 2013) on *D. simulans* and *D. mauritiana* EADs at 96 and 120 h AEL. We mapped our datasets against their reference genomes and also against the other species genome (see methods) to detect common, differentially accessible and species-specific regulatory regions (Fig. 4A). The ATAC-seq peak calling of the four datasets (two developmental stages and two species) revealed a total of 22 APREs (Associated Putative Regulatory Regions) in the *otd* locus, all of which were located within alignable orthologous regions in the two species (Fig. 4A).

Five of these peaks showed significant differences in accessibility between *D. mauritiana* and *D. simulans* at 96 h and/or 120 h: APRE 6 (*D. sim* chrX: 8,100,587-8,100,808, padj = 0.00155; *D. mau* chrX: 7,929,372-7,929,595, padj = 0.000877), APRE 7 (*D. mau* chrX: 7,929,910-7,930,230, padj = 1,85E-06) and APRE 11 (*D. sim* chrX: 8,107,881-8,108,402, padj = 0.00976; *D. mau* chrX: 7,936,681-7,937,179, padj = 0.0143) in the 3rd and 1st introns of *otd,* respectively, and APREs 17 (*D. sim* chrX: 8,125,765 - 8,126,100, padj = 0.0418; *D. mau* chrX: 7,954,305-7,954,661, padj = 0.0331) and 19 (*D. sim* chrX: 8,129,876 - 8,131,170, padj = 0.0317) located upstream of *otd* (Fig. 4A).

To test the activity of these APREs in L3 EADs, we generated *D. melanogaster* lines containing the APRE 6, 7-8, 11, 16-17, 19 sequences from *D. mauritiana* followed by GAL4 and crossed them to UAS-GFP (see Methods, Fig. 4B, Fig. S6). APREs 6 and 11 showed no activity in the EADs, which we confirmed using reporters with the equivalent *D. simulans* sequences (Fig. S6. APREs 16-17 and 19 drove GFP expression anterior to the MF, in the ocellar domain and some regions of the antennal disc (Fig S6). Finally, APRE7-8 drove expression in the posterior of the eye disc, in maturing ommatidia, in a similar pattern to endogenous *otd* expression, and consistent with APRE 7 and APRE 8 demarcating the 2.5 kb region corresponding to the *D. melanogaster otd^uvi^* allele previously identified as an *otd* eye enhancer (Vandendries et al., 1996) (Fig. 4A, Fig S7). We therefore compared the activity of the *D. mauritiana* and *D. simulans* APRE 7-8 sequences, and we observed differences in the onset of activity of these reporter lines. According to mRNA and protein Otd expression, *APRE7-8-GAL4* from *D. mauritiana* activated GFP expression earlier than the reporter with the *D. simulans* sequence (Fig. 4B-4D) consistent with the difference in endogenous *otd* expression between these species. These results suggest that differences in the *D. mauritiana* and *D. simulans* APRE 7-8 sequences underlie the difference in *otd* eye expression between these two species. Note that we also tested the activity of the *D. melanogaster* APRE 7-8 sequence and observed it was similar to the *D. simulans* sequence rather than that of *D. mauritiana* consistent with the derived larger ommatidia of the latter species (Fig S7).

We aligned the orthologous sequences of the 2.5 kb APRE 7-8 region from different *D. mauritiana, D. simulans* and *D. melanogaster* strains and found seventeen potentially fixed differences (thirteen SNPs and four short indels) in *D. mauritiana* Figs S8, S9). Interestingly most of these mutations lie in a 1.5 kb fragment of this enhancer identified by Vandendries et al (1996) and several fall in predicted binding sites for Sine oculis, Cut and Otd itself (as well as other transcription factors) (Fig. S9).

### Differences in Otd targets during eye development between *D. simulans* and *D. mauritiana*

Next, we investigated whether differences in the onset of expression of *otd* between *D. simulans* and *D. mauritiana* promoted further changes in the expression of downstream genes. To this end, we searched for the Otd-binding motif in accessible chromatin regions of genes expressed during eye development. Based on this analysis, we found 1,148 putative Otd target genes. We next examined which of these accessible chromatin regions were associated with genes that were differentially expressed in our transcriptome datasets. We found that 161 APREs of the 1330 genes that are upregulated in *D. mauritiana* contained Otd binding motifs, and 111 out of 1249 genes upregulated in *D. simulans* had associated peaks containing Otd binding motifs (Fig. 5A, C, Data S6).

**Figure 5.**
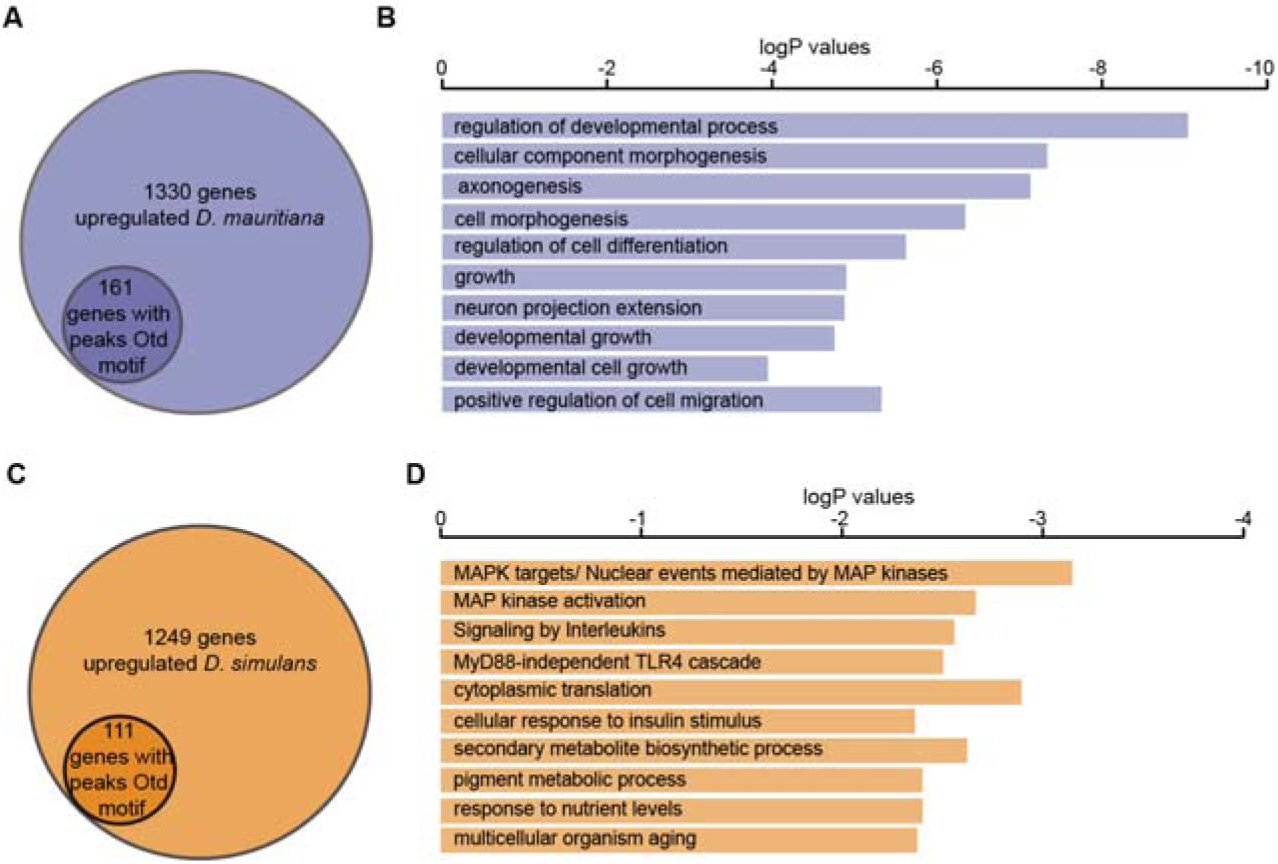
Otd downstream targets. **(A)** 161 genes upregulated in *D. mauritiana* have an associated peak that contains at least one Otd motif. **(B)** GO enrichment for those genes with Otd motifs in *D. mauritiana* **(C)** 111 genes upregulated in *D. simulans* have an associated peak that contains Otd motif. **(D)** GO enrichment for those genes with Otd motifs in *D. simulans*.

We then performed Gene Ontology (GO) term enrichment analysis for these differentially expressed genes with accessible chromatin containing Otd binding motifs. The *D. mauritiana* dataset exhibited enrichment in terms related to developmental processes, cell morphogenesis, axonogenesis, regulation of differentiation or growth, among others (Fig. 5B). By contrast, genes that were upregulated in *D. simulans* with associated Otd-peaks were enriched in terms such as the MAP kinase network, signalling by interleukins and cellular response to insulin stimulus (Fig. 5D).

## Discussion

While much is known about the specification and differentiation of ommatidia, very little is known about the regulation and evolution of their size (although see (Weinkove et al., 1999)). To investigate the genetic basis of the difference in ommatidia size between *D. mauritiana* and *D. simulans*, we carried out high-resolution introgression mapping of a previously identified X-linked QTL that explains about 33% of the difference in eye size between these two species (Arif et al., 2013a). In this region we identified eight positional candidate genes whose expression in the developing eye-antennal discs differed between *D. mauritiana* and *D. simulans*. Our analysis of the spatial expression of these eight genes strongly suggests *otd* as the best candidate gene in this region for contributing to the difference in ommatidia size and thus overall eye size between these species.

*otd/Otx* genes play several important roles during eye development in both invertebrates and vertebrates (Ragge et al., 2005; Ranade et al., 2008; Sen et al., 2013; Tahayato et al., 2003; Vandendries et al., 1996). During *Drosophila* eye development, Otd regulates genes for cell adhesion and cytoskeletal organisation, which is essential for the correct development of the photoreceptor cells and ommatidia maturation as well as subtype specification through regulation of rhodopsin expression (Fichelson et al., 2012; Johnston et al., 2011; Ranade et al., 2008; Tahayato et al., 2003). Mutations in *otd* perturb morphogenesis of the photoreceptor cells (Fichelson et al., 2012; Vandendries et al., 1996). Intriguingly, the removal of photoreceptor cells changes ommatidia size (Miller and Cagan, 1998). We propose that although *otd* is not expressed in the lens-secreting cone cells, it indirectly affects the organisation of these cells and thus ommatidia size through regulating the maturation and organisation of the underlying photoreceptor cells. We have shown that knockdown or loss of *otd* in *D. melanogaster* perturbs ommatidia size specification, but it remains to be directly tested if variation in the expression of this gene underlies larger and smaller ommatidia in *D. mauritiana* and *D. simulans* respectively and if *otd* contributes to the observed variation in ommatidia size in different regions of *Drosophila* eyes (Buffry et al., 2024; Gaspar et al., 2020).

Changes in developmental timing, or heterochrony, have played an essential role in the evolution of morphologies in multiple taxa (Alberch, P. et al., 1979; Alberch and Alberch, 1981; McKinney, 1988). Classically, the term heterochrony has been used to refer to differences in the timing of developmental events and several examples of heterochrony have been described (Briscoe and Small, 2015; Ebisuya and Briscoe, 2018; Farnworth et al., 2020; Keyte and Smith, 2014). Most of these characterised cases showed that the genetic basis lies upstream of the mechanism responsible for the heterochrony, such as changes in proliferating rates, differences in the initial size of the primordium or distinct rates of protein stability and biochemistry (Gomez et al., 2008; Kicheva et al., 2014; Matsuda et al., 2020; Rayon et al., 2020). Heterochronic shifts can also occur as direct consequence of the causative genetic change, such as those that affect regulatory regions altering the timing of gene expression (Ramaekers et al., 2019). Although differences in gene expression of single transcription factors have the potential to completely modify the subsequent GRN, the relative contribution of such direct heterochrony in generating morphological diversity remains unknown. Our data indicate that *otd* is actually expressed earlier during ommatidial maturation in *D. mauritiana* compared to *D. simulans.* This suggests that cis-regulatory changes in *otd* lead to ommatidial cells being exposed to Otd for longer in *D. mauritiana* resulting in larger ommatidia. Together with Ramaekers and colleagues (2019), our study shows how morphological diversity in closely related species may be achieved by subtly altering the temporal expression of a single transcription factor. Importantly, in both cases, these transcription factors, Ey and Otd, act upstream in the GRN controlling the developmental process, thus changes in their expression may promote major differences in downstream effectors.

Further exploration and comparison of the regulatory landscape of *otd* between *D. mauritiana* than *D. simulans* allowed us to identify an intronic region in the *otd* locus, APRE 7-8, corresponding to the orthologous sequence of the previously known “eye enhancer” from *D. melanogaster* (Bernardo-Garcia et al., 2016; Hauck et al., 1999; Vandendries et al., 1996). The *D. mauritiana* sequence reporter line showed earlier activity than *D. simulans* sequence, suggesting that this region is responsible for the differences in the timing of *otd* expression and therefore, function. Further work is needed to identify the transcription factors that directly bind to this enhancer and the changes in *D. mauritiana* sequence that underlie the differential activity of this element in this species (Fig. S8, Fig. S9).

We also investigated how these changes in *otd* expression might alter target gene expression to change ommatidia size. We identified a set of genes that are differentially expressed between these two species when the ommatidia are acquiring their final size that may be acting downstream of Otd, as they have accessible chromatin regions containing putative Otd binding motifs. We compared this set of genes to known and putative targets of Otd which have been characterised later in eye development during pupal stages (Fichelson et al., 2012; Ranade et al., 2008). This comparison showed that a subset of genes with altered expression in *otd* mutants are also differentially expressed between *D. mauritiana* and *D. simulans* in late L3. Among other TFs (*Dve, vnd, MED10*) we identified several genes involved in phototransduction (e.g. *slo, Slob, ninaG, inaD, ninaA*) and genes encoding cytoskeleton and adhesion proteins (*Act88F*) (Data S6). This further suggests that the network downstream of Otd varies between these two species and that ultimately, these changes in the GRN promote differences in ommatidia size between *D. mauritiana* and *D. simulans*.

Interestingly we recently showed that *D. mauritiana* has higher contrast sensitivity than *D. simulans* (Buffry et al., 2024), while the latter species has greater spatial acuity consistent with the differences in ommatidia size between these species. The trade-off between contrast sensitivity and acuity is heavily influenced by various aspects of visual ecology, such as habitat type, circadian activity patterns, and lifestyle. Thus, substantial functional consequences with strong ecological implications could be linked to changes in the expression of individual genes such as *otd*.

## Conclusions

Our data suggest that changes in the timing of *otd* expression underlie differences in ommatidia size and thus overall eye size between *D. mauritiana* and *D. simulans*. Our work provides new insights into ommatidia size regulation and the evolution of eye size. Together with evidence from other studies showing that changes in the timing of *ey* expression contributes to differences in ommatidia number in *Drosophila* (Ramaekers et al., 2019), we now have a better understanding of the genetic and developmental mechanisms that underlie the large diversity in *Drosophila* eye size (Arif et al., 2013a; Buchberger et al., 2021; Buffry et al., 2024; Gaspar et al., 2020; Hilbrant et al., 2014; Keesey et al., 2019; Norry and Gomez, 2017; Posnien et al., 2012; Ramaekers et al., 2019; Reis et al., 2020). Moreover, this evidence suggests that changes in the temporal expression of upstream transcription factors could be a widespread mechanism for morphological evolution.

## Materials and Methods

### Fly stocks and clonal analysis

*D. simulans* strain *yellow* (*y*), *vermillion* (*v*), *forked* (*f*) was obtained from the *Drosophila* Species Stock Center, San Diego, California (Stock no.14021–0251.146). *D. mauritiana* TAM16 is a wild-type inbred strain (Arif et al., 2013a). *UAS-miR-otd* and *UAS-otd* (III) were kindly provided by Henry Sun (Wang et al., 2010). *GMR-GAL4* (Hay et al., 1994) was used to drive expression of the transgenes. To generate mitotic clones of mutant *otd* in developing eyes, we used *otd[YH13], neoFRT19A/FM7c* and *RFP, neoFRT19A; ey-Flp*, which were obtained from Bloomington Stock Centre (#8675 and #67173 respectively).

*otd* mutant clones were induced in developing eyes using the Flp/FRT system. Female flies of the genotype *otd[YH13], neoFRT19A/FM7c* were crossed with males of the genotype *RFP, neoFRT19A; ey-Flp*. Female F1 progeny were examined for the lack of the Fm7c balancer and these flies were prepared for SEM analysis.

### Synchrotron radiation microtomography

Fly heads were removed from the body and placed into fixative (2% PFA, 2.5% GA in 0.1 M sodium cacodylate buffer over night at 4°C. Heads were washed in water, then placed into 1% osmium tetroxide for 48 hours at 4°C, then washed and dehydrated in increasing concentrations of ethanol up to 100%. Heads were then infiltrated with increasing 812 Epon resin concentrations up to 100 % over 5 days and polymerised in embedding moulds for 24 hrs at 70°C.

Heads were scanned at the TOMCAT beamline of the Swiss Light Source (Paul Scherrer Institute, Switzerland (Stampanoni et al., 2006). Scans were performed using a 16 keV monochromatic beam with a 20 µm LuAG:Ce scintillator. Resin blocks were trimmed and mounted using soft wax and scanned using 20x combined magnification (effective pixel size 325 nm) and a propagation distance of 25 mm. Two thousand projections were taken as the heads rotated through 180°, each with 200 ms exposure. Projections were reconstructed into 8-bit tiff stacks and Paganin filtered (delta = 1^-8^, beta = 2^-9^ (Paganin et al., 2002) using custom in-house software (Marone and Stampanoni, 2012). Tiff stacks were segmented in Amira (v2019.2, Thermo Fisher) for measurements of facet area.

### SEM microscopy

Fly heads were fixed in Bouin’s for 2 hours. Then, 1/3 of total volume was replaced by 100% ethanol to fully immerse heads in Bouin’s and were left to fix overnight. Heads were washed and dehydrated 2x 70% ethanol overnight, 2x in 100% ethanol and finally critical point dried and mounted onto sticky carbon tabs on SEM stubs, gold coated and imaged in a Hitachi S-3400N SEM with secondary electrons at 5kV.

### Markers and Introgression lines

Males were collected at backcross 7 of three replicate introgression lines (IL1, IL3 and IL4) that were recombinant within the introgressed region (males with phenotypes: *y, f* or *v, f*). These individuals were genotyped with eleven new additional markers (Data S2).

Significant association between each marker and eye size was tested (F-test, type III sum of squares SS) by performing a single-marker ANOVA on the residuals of eye area regressed onto T1 tibia length for each replicate (introgression line (IL 1,3 and 4; n = 20 – 60, Data S2). Multiple testing was corrected using Bonferroni correction. All ANOVA models were fitted in the R statistical environment (R Development Core Team 2012) using the CAR package (Fox and Weisberg, 2019).

To narrow down the 2 Mb region, the X chromosome region between *y* and *v* from *D. mauritiana* TAM16 was re-introgressed into *D. simulans y, v, f* (as in Arif et al. 2013) and *y, f* females were backcrossed from multiple replicate lines to *y, v, f* males for a further nine generations. At the end of the egg-laying cycle of that generation, we collected mothers and genotyped them for molecular markers located in the 2 Mb region (Data S2). Four mothers with breakpoints within this region were identified. Two of them were siblings (IL9.1a and IL9.1b) and they had the same 4^th^ great-grandmother as IL9.3 and the same 7^th^ great-grandmother as IL9.2. Male progeny available for each of these females was collected and genotyped and phenotyped for eye area, ommatidia diameter, ommatidia number and T1 tibia length as described previously in Posnien et al. (2012) (Data S2). To determine if the *D. mauritiana* DNA in the 2 Mb region resulted in larger eyes and larger ommatidia, *y*, *f* males (i.e. with some *D. mauritiana* DNA in the 2 Mb interval) were compared to that of their *y, v, f* sibling males (i.e. without *D. mauritiana* DNA) for each introgression line using one-tailed, two-sample, equal-variance t-tests.

### In situ hybridisation and immunohistochemistry

In situ hybridizations were carried out using a standard protocol with DIG-labelled antisense RNA probes. Eye-antenna imaginal discs were dissected and fixed at 120 h AEL for 40 min in 4% formaldehyde. To be able to compare the expression patterns avoiding technical differences (i.e. probe affinity and probe concentration), we first aligned the sequences from *D. mauritiana* and *D. simulans* and designed RNA probes within fragments with at least 95% of similarity between them (Data S8). This design allowed us to perform the *in situ* hybridization experiments using the same specific probes for each of the candidate genes at the same concentration for both species. The nitro blue tetrazolium/5-bromo-4-chloro-3′-indolyphosphate (NBT-BCIP) reaction was stopped at the same time. Candidate gene sequences were cloned into a TOPO PCR4 (*spirit, otd*, *Ppt1, CG1632, Es2* and *CG12112*) or pCRII (*CG1885, Sptr*) vectors (Invitrogen) using specific primer pairs (Data S8), respectively, following the manufacturer’s protocol. M13 forward and reverse primers were used to linearize the DNA. According to the vector and orientation of the fragments T3, T7 or SP6 RNA polymerase were used to generate the DIG-labelled riboprobes.

Immunostainings with Rabbit anti□Otd (Wang et al. 2010) and rat anti-Elav (7E8A10, Hybridoma bank) were performed at 1:1500 and 1:100 dilutions respectively using standard protocols, followed by anti□rat□Cy3 (Jackson Immuno Research) and anti-rabbit-Alexafluor 647 (Molecular probes) secondary AB staining, at 1:200. The actin cytoskeleton was stained with Alexafluor 488□Phalloidin (Molecular Probes) at 1:40 dilution for 30□min after secondary antibody incubation. Discs were mounted in Prolong Gold antifade reagent, supplemented with DAPI (Molecular Probes), and captured with a Zeiss LSM 510 confocal microscope. Images were processed using NIH ImageJ software.

### RNA-seq

Flies were raised at 25°C with a 12h:12h dark:light cycle and their eggs were collected in 2 h time periods. Freshly hatched L1 larvae were transferred into fresh vials in density-controlled conditions (30 freshly hatched L1 larvae per vial). EADs were dissected at three different developmental time points: 72 96 and 120 h AEL and stored in RNALater (Qiagen, Venlo, Netherlands). Three biological replicates for each sample were generated. Total RNA was isolated using RNeasy Mini Kit (Qiagen). RNA quality was determined using the Agilent 2100 Bioanalyzer (Agilent Technologies, Santa Clara, CA, USA) microfluidic electrophoresis.

Library preparation for RNA-seq was performed using the TruSeq RNA Sample Preparation Kit (Illumina, catalog ID RS-122-2002) starting from 500 ng of total RNA. Accurate quantitation of cDNA libraries was performed using the QuantiFluor™dsDNA System (Promega, Madison, Wisconsin, USA). The size range of final cDNA libraries was determined by applying the DNA 1000 chip on the Bioanalyzer 2100 from Agilent (280 bp). cDNA libraries were amplified and sequenced using cBot and HiSeq 2000 (Illumina): only 120 h EAD samples were sequenced as paired-end (PE) reads (2 x 100 bp), all the other samples were sequenced in single-end (SE) reads (1 x 50 bp). Sequence images were transformed to bcl files using the software BaseCaller (Illumina). The bcl files were demultiplexed to fastq files with CASAVA (version 1.8.2).

Quality control analysis using FastQC software (version 0.10.1, Babraham Bioinformatics) was performed. All RNAseq reads are accessible in the Short Read Archive through umbrella BioProject PRJNA666691 (containing PRJNA374838 and PRJNA666524). Before the mapping step, PE 100 bp reads were converted into SE 50 bp by splitting the reads in half and merging right and left reads into a single file.

The reciprocally re-annotated references described in (Torres-Oliva et al., 2016) were used to map the species-specific reads. Bowtie2 (Langmead and Salzberg, 2012) was used to map the reads to each reference (–very-sensitive-local –N 1) and the idxstats command from SAMtools v0.1.19 (Li et al., 2009) was used to summarize the number of mapped reads. HTSFilter (Rau et al., 2013) was used with default parameters to filter out genes with very low expression in all samples. For the remaining genes in each pair-wise comparison, differential expression was calculated using DESeq2 v1.2.7. with default parameters (Love et al., 2014).

### ATAC-seq library preparation and sequencing

Samples were obtained following the same procedure as for the RNA-seq experiments: flies were raised at 25° C with a 12h:12h dark:light cycle. Freshly hatched L1 larvae were transferred into vials with density-controlled conditions. EADs were dissected at 96 and 120 h AEL and maintained in ice cold PBS. Imaginal disc cells were lysed in 50 μl Lysis Buffer (10 mM Tris-HCl, pH = 7.5; 10 mM NaCl; 3 mM MgCl_2_; 0.1% IGEPAL). Nuclei were collected by centrifugation at 500 g for 5 min. 75,000 nuclei were suspended in 50 μl Tagmentation Mix [25 μl Buffer (20 mM Tris-CH_3_COO^-^, pH = 7.6; 10 mM MgCl_2_; 20% Dimethylformamide); 2.5 μl Tn5 Transposase; 22.5 μl H_2_O] and incubated at 37 LJC for 30 min. After addition of 3 μl 2 M NaAC, pH = 5.2 DNA was purified using a QIAGEN MinElute Kit. PCR amplification for library preparation was done for 14 cycles with NEBNext High Fidelity Kit; primers were used according to (Buenrostro et al., 2013). Paired end 50 bp sequencing was carried out by the Transcriptome and Genome Analysis Laboratory Goettingen, Germany.

### ATAC-seq peak calling and differential binding site analysis

ATAC-seq raw reads were generated from the following samples (2 replicates each): *D. simulans* larvae at 96 and 120 h AEL and *D. mauritiana* larvae at 96 and 120 h AEL. These reads were mapped to strain-specific genomes of *D. mauritiana* and *D. simulans* (Torres-Oliva et al., 2016) using Bowtie2 (version 2.3.4.1) (Langmead and Salzberg, 2012) with the parameter –X2000. The Samtools suite v0.1.19 (Li et al., 2009) was used to convert *.sam to *.bam files and to further process the mapped reads. Duplicates were removed using Picard (version 2.20.2) with the parameter REMOVE_DUPLICATE=TRUE. Bam files were then converted to bed files using the Bedtools (version 2.24) bamtobed command. Reads were centred according to (Buenrostro et al., 2015). These reads were then converted to the *D. simulans* or *D. mauritiana* coordinate system using liftOver (1.14.0) with custom prepared chain files, one for the conversion of *D. mauritiana* coordinates to *D. simulans* coordinates and one for the conversion of *D. simulans* coordinates to *D. mauritiana* coordinates. Peaks were then called using MACS2 (version 2.1.2, (Zhang et al., 2008)) with the following parameters: --shift – 100, extsize 200, -q 0.01.

We used the Diffbind package (version 2.12.0, (Ross-Innes et al., 2012)) in R (version 3.6.1.) to search for differentially accessible ATAC-seq regions. A consensus peak set of 19,872 peaks (96 h AEL) and 15,868 peaks (120 h AEL) was used for all samples and the reads were counted for each identified peak with the dba.count command. For each time point separately we used the dba.analyze command with default parameters to get differentially accessible peaks between the two species. This command uses by default the DESeq2 analysis. All plots were generated with the DiffBind package.

### Reporter assays

All APREs with the exception of *D. mauritiana-APRE6, D. simulans-APRE11 and D. mauritiana-APRE11,* were cloned upstream of GAL4 into the pBPGUw backbone (a gift from Gerald Rubin, Addgene plasmid 17575) by Genewiz. *D. mauritiana-APRE6, D. simulans-APRE11 and D. mauritiana-APRE11* were first sub-cloned into the pENTR-TOPO-D plasmid backbone then shuttled into the pBPGUw backbone using Gateway cloning. Sequences of all APREs used can be found in Data S9.

*otd-APRE7-8-GAL4* lines with *D. mauritiana* and *D. simulans* sequences were crossed with *UAS-nlsGFP* to characterize the activity of APRE7-8. EADs were dissected and fixed according to standard protocols. Immunostaining was performed with mouse anti-Elav (Hybridoma Bank) at 1:100 and rabbit anti-GFP (Molecular probes) at 1:1000, followed by secondary antibody incubation with anti-rabbit Alexa Fluor 488 and anti-mouse Alexa Fluor 568, both at 1:400. Imaging was performed using a Leica Stellaris microscope with a 40X objective and zoom 2. Images were processed using NIH ImageJ software.

We used Elav staining to count the number of ommatidial rows as a measure of developmental stage. We define the first row showing full enhancer activity as the first row counting from the posterior margin of the disc with more than 90% of the ommatidia positive for enhancer signal/GFP signal. For all discs, we determined the developmental time and the first row with full enhancer activity. Then we calculated the distance between the morphogenetic furrow and the first row of full enhancer activity as the difference between the total ommatidial rows and the first ommatidial row with full enhancer activity.

We then plotted the first row with full enhancer activity at different developmental times and fitted a regression line to model the observed linear trend for each species, with calculated R value for correlation and p value of fit.

Violin plots were used to show the distribution of distances between the morphogenetic furrow and the first row of full enhancer activity. We used Welch’s t-test to test whether the difference in distances between species was significant. In the case of the three-species comparison (Fig. S7), we use the nonparametric Games-Howell test to make pairwise comparisons among the three species. We used an Analysis of Covariance (ANCOVA) to test for differences between strains for Otd-positive ommatidia while adjusting for differences in development stage by using the number of ommatidial rows as a proxy for the latter. The ANCOVA was performed using base R v4.0.2 (R Core Team, 2020).

### Prediction of transcription factor binding sites and potential target genes

To search for TFBM of potential *otd* regulators, we used the universalmotif package (version 1.20.0) using its function to scan sequences. The JASPAR core and CISBP databases were used for screening DNA sequences of ATAC-seq peaks in the *otd* APRE7-8 sequence with all possible TFBSs from *Drosophila* with a threshold of p-value >0.001. Potential fixed changes in the 2.5 kb APRE7-8 sequence of *D. mauritiana* were inferred from aligning the sequences of this region from strains of *D. mauritiana* (Red3, mau12, mav2 and TAM16), *D. simulans* (m3 and w501) and the *D. melanogaster* sequence using Clustal Omega with default parameters (Li et al., 2009)(Fig. S9).

To define a list of potential Otd target genes, we used an Otd-motif (Dmelanogaster-FlyFactorSurvey-Oc_Cell_FBgn0004102) from the MotifDB package (version 1.16.1), which provides a collection of available transcription factors in R (version 3.3.3). We searched for Otd binding sites in accessible chromatin regions with the findMotifsGenome.pl command implemented in the HOMER (version V4.10.4, (Heinz et al., 2010)) in all samples. All peaks with a predicted Otd motif were annotated to an associated gene using the annotatePeaks.pl command by HOMER and combined all time points and both species into one file. We then looked for the number of genes with an annotated Otd motif and found 1,148 unique genes, which we overlapped with our RNA-seq dataset to find out which of these target genes were differentially expressed between the two species. GO term enrichment analysis of putative Otd target genes was performed using the online tool Metascape (Zhou et al., 2019).

We used the online STRING database that integrates all known and predicted associations between proteins based on evidence from a variety of sources (Szklarczyk et al., 2021), to construct networks of DEG encoded proteins. To visualize the network and map genes/prot with Otd motifs we applied the Cytoscape software (Shannon et al., 2003).

## Data Availability

All RNAseq and ATACseq reads are accessible in the Short Read Archive through umbrella BioProject PRJNA666691 (containing PRJNA374838 and PRJNA666524).

## Acknowledgements

This work was funded by grants from the ERC (242553) to A.P.M. and BBSRC (BB/M020967/1, BB/T000317/1) to A.P.M. and I.A., and A.P.M. and M.K respectively. M.T.-O. was supported by a “la Caixa-DAAD” fellowship and and GG by MICINN contract PRE2019-090291. N.P. and E.B. were funded by the Deutsche Forschungsgesellschaft (DFG, Grant No. PO 1648/3-1 to N.P.). We acknowledge the Paul Scherrer Institut, Villigen, Switzerland for provision of synchrotron radiation beamtime at the TOMCAT beamline X02DA of the SLS, and the invaluable assistance of Dr Christian Schlepütz. We thank the Transcriptome and Genome Core Unit, University Medical Center Göttingen (UMG) for support with next generation sequencing, the Advanced Light Microscopy and Image Analysis (ALMIA) platform, at CABD, for imaging support, and Franck Pichaud (UCL, UK) for access to *otd* microarray data. We also thank Madeleine Aase-Remedios and Sebastian Kittelmann for assistance with sequence analysis.

## Author contributions

M.T.-O. performed and analysed RNA-seq experiments, interpreted the results and prepared figures. E.B. Bioinformatics analysis and figures. A.D.B. performed functional tests of otd perturbations and reporter studies, interpreted the results. M.K. Synchrotron radiation microtomography and interpretation of the data G.G reporter studies and interpretation. L.S.-R. Synchrotron radiation microtomography. P.G. gene expression experiments G.C.B. gene expression experiments. J.F.J. enhancer sequence analysis. F.C. enhancer data interpretation and supervision. S.A. and M.D.S.N. fine-mapping and introgression. N.P. supervision, funding and data analysis interpretation. A.P.M. and I.A. work conceptualization, supervision, design, data analysis and interpretation, figures, funding and manuscript writing with the help of all other authors.

## Supplementary Files

**Figure S1. RNA-seq datasets. (A)** Heat-map of all RNA-seq samples. **(B)** PCA plot of all RNA-seq samples.

**Figure S2. *otd* expression in pupal eyes. (A)** *D. mauritiana* pupal eye (48 h APF) stained with Phalloidin (Actin, A’), anti-Elav marks photoreceptors (A’’) and Otd is expressed in all ommatidia (A’’’). **(B)** Otd is present in all ommatidia in *D. simulans* pupal eyes, (b’) Actin (Phalloidin) highlights ommatidia area, (B’’) anti-Elav marks photoreceptors and Otd protein is shown in B’’’.

**Figure S3. Loss of *otd* causes defects in ommatidia structure. (A-C)** Knockdown of *otd* by expressing *UAS-miR-otd* with *GMR-GAL4* driver in *D. melanogaster* eye. (a-a’) *GMR-GAL.* **(B-B’)** *GMR-GAL/ UAS-miR-otd* shows a rough eye phenotype due to defects in ommatidia arrangements. **(C-C’)** *GMR-GAL/ UAS-miR-otd* phenotype is rescued by co-expressing *UAS-otd*. **(D-D’)** *otd* mutant mitotic clones also show defects in ommatidia size and organisation.

**Figure S4. Alignment of *D. mauritiana* TAM16 and *D. simulans y*, *v*, *f* Otd protein sequences**

**Figure S5. Topological Domain Associated with *otd* locus in *D. melanogaster*** from http://chorogenome.ie-freiburg.mpg.de/

**Figure S6. Activity of tested APREs in the otd locus of D. simulans and D. mauritiana.** L3 instar eye-antenna imaginal discs from *D. melanogaster* lines containing *D. mauritiana-APRE4-GAL4, D. mauritiana-APRE6-GAL4, D. mauritiana-APRE7-8-GAL4, D. simulans-APRE11-GAL4, D. mauritiana-APRE13-14-GAL4, D. mauritiana-APRE16-17-GAL4, D. mauritiana-APRE19-GAL4* crossed to *UAS-GFP* and stained with DAPI (grey). Scale bar = 50μm.

**Figure S7. *D. melanogaster, D. simulans* and *D. mauritiana otd-APRE7-8* enhancer activity in 3^rd^ instar larvae eye imaginal discs. (A)** *D. melanogaster, D. simulans* and *D. mauritiana otd-APRE7-8* 3^rd^ instar larvae eye imaginal discs immunostained with anti-Elav antibody to visualise the progression of ommatidia maturation as proxy for the developmental time. **(B)** Plot showing the number of GFP-positive ommatidia rows (x-axis) activated by *otd-APRE7-8* at different developmental time points (y-axis, developmental points inferred by number of ommatidia rows) for both species. **(C)** Violin plots showing the distance between MF and the first row of *otd-APRE7-8* activity. *D. mauritiana otd-APRE7-8* is active earlier during the differentiation of ommatidia.

**Figure S8.** Clustal Omega alignment of the 2.5 kb APRE7-8 region from selected strains of *D. mauritiana*, *D. simulans* and *D. melanogaster* including the sequences used in enhancer reporter constructs for each species (*otd-APRE7-8*).

**Figure S9.** Schematic showing the position of the 2.5 kb APRE7-8 within the *otd* locus on the X chromosome of *D. melanogaster*. A 1.5 kb region shown to drive similar expression to the 2.6 kb region is indicated by a purple bar (Vandendries et al., 1996). Below is shown the ATAC-seq profile for this region and the positions of potential fixed mutations specific to *D. mauritiana* (arrows indicate SNPs and rectangles indicate short indels) with predicted binding sites for Otd, So and Cut in *D. melanogaster* indicated by arrows. Binding sites indicated with an asterisk contain mutations in *D. mauritiana*.

**Data S1.** Measurements of ommatidia size of *D. mauritiana* and *D. simulans* eyes.

**Data S2.** Mapping and introgression data.

**Data S3.** Pair-wise differential gene expression analysis (D.sim vs. D.mau) results for each time point (72h, 96h and 120h).

**Data S4.** Measurements of Otd-positive ommatidia in *D. mauritiana* and *D. simulans* eye discs

**Data S5.** Differential peak calling for *D. mau* and *D. sim* 96 and 120 h AEL ATAC-seq data

**Data S6.** TFBM in differential open chromatin regions in *otd* locus of *D. simulans* and *D. mauritiana*

**Data S7.** Differentially expressed genes with predicted Otd TFBMs in their associated open chromatin regions.

**Data S8.** Primers used to generate probes for in situ hybridization.

**Data S9.** Sequences used to generate reporter lines.

## References

Alberch, P., Alberch, J., 1981. Heterochronic mechanisms of morphological diversification and evolutionary change in the neotropical salamander, Bolitoglossa occidentalis (Amphibia: Plethodontidae). J. Morphol. 167, 249–264. 10.1002/jmor.1051670208

Alberch, P., Gould, S.J., Oster, G.F, Wake, D.B, 1979. Size and Shape in Ontogeny and Phylogeny. Paleobiology 5, 296–317.

Arif, S., Hilbrant, M., Hopfen, C., Almudi, I., Nunes, M.D.S., Posnien, N., Kuncheria, L., Tanaka, K., Mitteroecker, P., Schlötterer, C., McGregor, A.P., 2013a. Genetic and developmental analysis of differences in eye and face morphology between Drosophila simulans and Drosophila mauritiana. Evol. Dev. 15, 257–267. 10.1111/ede.12027

Arif, S., Murat, S., Almudi, I., Nunes, M.D.S., Bortolamiol-Becet, D., McGregor, N.S., Currie, J.M.S., Hughes, H., Ronshaugen, M., Sucena, É., Lai, E.C., Schlötterer, C., McGregor, A.P., 2013b. Evolution of mir-92a underlies natural morphological variation in Drosophila melanogaster. Curr. Biol. CB 23, 523–528. 10.1016/j.cub.2013.02.018

Arnoult, L., Su, K.F.Y., Manoel, D., Minervino, C., Magriña, J., Gompel, N., Prud’homme, B., 2013. Emergence and diversification of fly pigmentation through evolution of a gene regulatory module. Science 339, 1423–1426. 10.1126/science.1233749

Bernardo-Garcia, F.J., Fritsch, C., Sprecher, S.G., 2016. The transcription factor Glass links eye field specification with photoreceptor differentiation in Drosophila. Dev. Camb. Engl. 143, 1413–1423. 10.1242/dev.128801

Briscoe, J., Small, S., 2015. Morphogen rules: design principles of gradient-mediated embryo patterning. Dev. Camb. Engl. 142, 3996–4009. 10.1242/dev.129452

Buchberger, E., Bilen, A., Ayaz, S., Salamanca, D., Matas de Las Heras, C., Niksic, A., Almudi, I., Torres-Oliva, M., Casares, F., Posnien, N., 2021. Variation in Pleiotropic Hub Gene Expression Is Associated with Interspecific Differences in Head Shape and Eye Size in Drosophila. Mol. Biol. Evol. 38, 1924–1942. 10.1093/molbev/msaa335

Buenrostro, J.D., Giresi, P.G., Zaba, L.C., Chang, H.Y., Greenleaf, W.J., 2013. Transposition of native chromatin for fast and sensitive epigenomic profiling of open chromatin, DNA-binding proteins and nucleosome position. Nat. Methods 10, 1213–1218. 10.1038/nmeth.2688

Buenrostro, J.D., Wu, B., Chang, H.Y., Greenleaf, W.J., 2015. ATACLseq: A Method for Assaying Chromatin Accessibility GenomeLWide. Curr. Protoc. Mol. Biol. 109. 10.1002/0471142727.mb2129s109

Buffry, A.D., Currea, J.P., Franke-Gerth, F.A., Palavalli-Nettimi, R., Bodey, A.J., Rau, C., Samadi, N., Gstöhl, S.J., Schlepütz, C.M., McGregor, A.P., Sumner-Rooney, L., Theobald, J., Kittelmann, M., 2024. Evolution of compound eye morphology underlies differences in vision between closely related Drosophila species. BMC Biol. 22, 67. 10.1186/s12915-024-01864-7

Casares, F., Almudi, I., 2016. Fast and Furious 800. The Retinal Determination Gene Network in Drosophila, in: Castelli-Gair Hombría, J., Bovolenta, P. (Eds.), Organogenetic Gene Networks. Springer International Publishing, Cham, pp. 95–124. 10.1007/978-3-319-42767-6_4

Casares, F., McGregor, A.P., 2021. The evolution and development of eye size in flies. Wiley Interdiscip. Rev. Dev. Biol. 10, e380. 10.1002/wdev.380

Courtier-Orgogozo, V., Arnoult, L., Prigent, S.R., Wiltgen, S., Martin, A., 2020. Gephebase, a database of genotype-phenotype relationships for natural and domesticated variation in Eukaryotes. Nucleic Acids Res. 48, D696–D703. 10.1093/nar/gkz796

Currea, J.P., Smith, J.L., Theobald, J.C., 2018. Small fruit flies sacrifice temporal acuity to maintain contrast sensitivity. Vision Res. 149, 1–8. 10.1016/j.visres.2018.05.007

Duncan, A.B., Salazar, B.A., Garcia, S.R., Brandley, N.C., 2021. A Sexual Dimorphism in the Spatial Vision of North American Band-Winged Grasshoppers. Integr. Org. Biol. Oxf. Engl. 3, obab008. 10.1093/iob/obab008

Ebisuya, M., Briscoe, J., 2018. What does time mean in development? Dev. Camb. Engl. 145, dev164368. 10.1242/dev.164368

Farnworth, M.S., Eckermann, K.N., Bucher, G., 2020. Sequence heterochrony led to a gain of functionality in an immature stage of the central complex: A fly–beetle insight. PLOS Biol. 18, e3000881. 10.1371/journal.pbio.3000881

Fichelson, P., Brigui, A., Pichaud, F., 2012. Orthodenticle and Kruppel homolog 1 regulate Drosophila photoreceptor maturation. Proc. Natl. Acad. Sci. U. S. A. 109, 7893–7898. 10.1073/pnas.1120276109

Fox, J., Weisberg, S., 2019. An R companion to applied regression, Third edition. ed. SAGE, Los Angeles.

Gaspar, P., Arif, S., Sumner-Rooney, L., Kittelmann, M., Bodey, A.J., Stern, D.L., Nunes, M.D.S., McGregor, A.P., 2020. Characterization of the Genetic Architecture Underlying Eye Size Variation Within Drosophila melanogaster and Drosophila simulans. G3 Bethesda Md 10, 1005–1018. 10.1534/g3.119.400877

Gomez, C., Ozbudak, E.M., Wunderlich, J., Baumann, D., Lewis, J., Pourquié, O., 2008. Control of segment number in vertebrate embryos. Nature 454, 335–339. 10.1038/nature07020

Gonzalez-Bellido, P.T., Wardill, T.J., Juusola, M., 2011. Compound eyes and retinal information processing in miniature dipteran species match their specific ecological demands. Proc. Natl. Acad. Sci. U. S. A. 108, 4224–4229. 10.1073/pnas.1014438108

Hauck, B., Gehring, W.J., Walldorf, U., 1999. Functional analysis of an eye specific enhancer of the eyeless gene in Drosophila. Proc. Natl. Acad. Sci. U. S. A. 96, 564–569. 10.1073/pnas.96.2.564

Hay, B.A., Wolff, T., Rubin, G.M., 1994. Expression of baculovirus P35 prevents cell death in Drosophila. Dev. Camb. Engl. 120, 2121–2129. 10.1242/dev.120.8.2121

Heinz, S., Benner, C., Spann, N., Bertolino, E., Lin, Y.C., Laslo, P., Cheng, J.X., Murre, C., Singh, H., Glass, C.K., 2010. Simple Combinations of Lineage-Determining Transcription Factors Prime cis-Regulatory Elements Required for Macrophage and B Cell Identities. Mol. Cell 38, 576–589. 10.1016/j.molcel.2010.05.004

Hilbrant, M., Almudi, I., Leite, D.J., Kuncheria, L., Posnien, N., Nunes, M.D.S., McGregor, A.P., 2014. Sexual dimorphism and natural variation within and among species in the Drosophila retinal mosaic. BMC Evol. Biol. 14, 240. 10.1186/s12862-014-0240-x

Horridge, G.A., 1977. The Compound Eye of Insects. Sci. Am. 237, 108–120. 10.1038/scientificamerican0777-108

Johnston, R.J., Otake, Y., Sood, P., Vogt, N., Behnia, R., Vasiliauskas, D., McDonald, E., Xie, B., Koenig, S., Wolf, R., Cook, T., Gebelein, B., Kussell, E., Nakagoshi, H., Desplan, C., 2011. Interlocked feedforward loops control cell-type-specific Rhodopsin expression in the Drosophila eye. Cell 145, 956–968. 10.1016/j.cell.2011.05.003

Keesey, I.W., Grabe, V., Gruber, L., Koerte, S., Obiero, G.F., Bolton, G., Khallaf, M.A., Kunert, G., Lavista-Llanos, S., Valenzano, D.R., Rybak, J., Barrett, B.A., Knaden, M., Hansson, B.S., 2019. Inverse resource allocation between vision and olfaction across the genus Drosophila. Nat. Commun. 10, 1162. 10.1038/s41467-019-09087-z

Keyte, A.L., Smith, K.K., 2014. Heterochrony and developmental timing mechanisms: changing ontogenies in evolution. Semin. Cell Dev. Biol. 34, 99–107. 10.1016/j.semcdb.2014.06.015

Kicheva, A., Bollenbach, T., Ribeiro, A., Valle, H.P., Lovell-Badge, R., Episkopou, V., Briscoe, J., 2014. Coordination of progenitor specification and growth in mouse and chick spinal cord. Science 345, 1254927. 10.1126/science.1254927

Kittelmann, M., McGregor, A.P., 2024. Looking across the gap: Understanding the evolution of eyes and vision among insects. BioEssays News Rev. Mol. Cell. Dev. Biol. 46, e2300240. 10.1002/bies.202300240

Klaassen, H., Wang, Y., Adamski, K., Rohner, N., Kowalko, J.E., 2018. CRISPR mutagenesis confirms the role of oca2 in melanin pigmentation in Astyanax mexicanus. Dev. Biol. 441, 313–318. 10.1016/j.ydbio.2018.03.014

Kumar, J.P., 2018. The fly eye: Through the looking glass. Dev. Dyn. Off. Publ. Am. Assoc. Anat. 247, 111–123. 10.1002/dvdy.24585

Land, M.F., 1997. Visual acuity in insects. Annu. Rev. Entomol. 42, 147–177. 10.1146/annurev.ento.42.1.147

Land, M.F., 1989. Variations in the Structure and Design of Compound Eyes, in: Stavenga, D.G., Hardie, R.C. (Eds.), Facets of Vision. Springer Berlin Heidelberg, Berlin, Heidelberg, pp. 90–111. 10.1007/978-3-642-74082-4_5

Land, Nilsson, 2012. Animal eyes, 2nd ed. ed, Oxford animal biology series. Oxford university press, Oxford.

Langmead, B., Salzberg, S.L., 2012. Fast gapped-read alignment with Bowtie 2. Nat. Methods 9, 357–359. 10.1038/nmeth.1923

Li, H., Handsaker, B., Wysoker, A., Fennell, T., Ruan, J., Homer, N., Marth, G., Abecasis, G., Durbin, R., 1000 Genome Project Data Processing Subgroup, 2009. The Sequence Alignment/Map format and SAMtools. Bioinforma. Oxf. Engl. 25, 2078–2079. 10.1093/bioinformatics/btp352

Love, M.I., Huber, W., Anders, S., 2014. Moderated estimation of fold change and dispersion for RNA-seq data with DESeq2. Genome Biol. 15, 550. 10.1186/s13059-014-0550-8

Marone, F., Stampanoni, M., 2012. Regridding reconstruction algorithm for real-time tomographic imaging. J. Synchrotron Radiat. 19, 1029–1037. 10.1107/S0909049512032864

Matsuda, M., Hayashi, H., Garcia-Ojalvo, J., Yoshioka-Kobayashi, K., Kageyama, R., Yamanaka, Y., Ikeya, M., Toguchida, J., Alev, C., Ebisuya, M., 2020. Species-specific segmentation clock periods are due to differential biochemical reaction speeds. Science 369, 1450–1455. 10.1126/science.aba7668

McKinney, M.L. (Ed.), 1988. Heterochrony in Evolution: A Multidisciplinary Approach, Topics in Geobiology. Springer US, Boston, MA. 10.1007/978-1-4899-0795-0

Mencarelli, C., Pichaud, F., 2015. Orthodenticle Is Required for the Expression of Principal Recognition Molecules That Control Axon Targeting in the Drosophila Retina. PLoS Genet. 11, e1005303. 10.1371/journal.pgen.1005303

Miller, D.T., Cagan, R.L., 1998. Local induction of patterning and programmed cell death in the developing Drosophila retina. Dev. Camb. Engl. 125, 2327–2335. 10.1242/dev.125.12.2327

Norry, F.M., Gomez, F.H., 2017. Quantitative Trait Loci and Antagonistic Associations for Two Developmentally Related Traits in the Drosophila Head. J. Insect Sci. Online 17, 19. 10.1093/jisesa/iew115

Paganin, D., Mayo, S.C., Gureyev, T.E., Miller, P.R., Wilkins, S.W., 2002. Simultaneous phase and amplitude extraction from a single defocused image of a homogeneous object. J. Microsc. 206, 33–40. 10.1046/j.1365-2818.2002.01010.x

Palavalli-Nettimi, R., Theobald, J.C., 2020. Small eyes in dim light: Implications to spatio-temporal visual abilities in Drosophila melanogaster. Vision Res. 169, 33–40. 10.1016/j.visres.2020.02.007

Perl, C.D., Niven, J.E., 2016. Differential scaling within an insect compound eye. Biol. Lett. 12, 20160042. 10.1098/rsbl.2016.0042

Posnien, N., Hopfen, C., Hilbrant, M., Ramos-Womack, M., Murat, S., Schönauer, A., Herbert, S.L., Nunes, M.D.S., Arif, S., Breuker, C.J., Schlötterer, C., Mitteroecker, P., McGregor, A.P., 2012. Evolution of eye morphology and rhodopsin expression in the Drosophila melanogaster species subgroup. PloS One 7, e37346. 10.1371/journal.pone.0037346

Ragge, N.K., Brown, A.G., Poloschek, C.M., Lorenz, B., Henderson, R.A., Clarke, M.P., Russell-Eggitt, I., Fielder, A., Gerrelli, D., Martinez-Barbera, J.P., Ruddle, P., Hurst, J., Collin, J.R.O., Salt, A., Cooper, S.T., Thompson, P.J., Sisodiya, S.M., Williamson, K.A., Fitzpatrick, D.R., van Heyningen, V., Hanson, I.M., 2005. Heterozygous mutations of OTX2 cause severe ocular malformations. Am. J. Hum. Genet. 76, 1008–1022. 10.1086/430721

Ramaekers, A., Claeys, A., Kapun, M., Mouchel-Vielh, E., Potier, D., Weinberger, S., Grillenzoni, N., Dardalhon-Cuménal, D., Yan, J., Wolf, R., Flatt, T., Buchner, E., Hassan, B.A., 2019. Altering the Temporal Regulation of One Transcription Factor Drives Evolutionary Trade-Offs between Head Sensory Organs. Dev. Cell 50, 780–792.e7. 10.1016/j.devcel.2019.07.027

Ranade, S.S., Yang-Zhou, D., Kong, S.W., McDonald, E.C., Cook, T.A., Pignoni, F., 2008. Analysis of the Otd-dependent transcriptome supports the evolutionary conservation of CRX/OTX/OTD functions in flies and vertebrates. Dev. Biol. 315, 521–534. 10.1016/j.ydbio.2007.12.017

Rau, A., Gallopin, M., Celeux, G., Jaffrézic, F., 2013. Data-based filtering for replicated high-throughput transcriptome sequencing experiments. Bioinforma. Oxf. Engl. 29, 2146– 2152. 10.1093/bioinformatics/btt350

Rayon, T., Stamataki, D., Perez-Carrasco, R., Garcia-Perez, L., Barrington, C., Melchionda, M., Exelby, K., Lazaro, J., Tybulewicz, V.L.J., Fisher, E.M.C., Briscoe, J., 2020. Species-specific pace of development is associated with differences in protein stability. Science 369, eaba7667. 10.1126/science.aba7667

Reis, M., Wiegleb, G., Claude, J., Lata, R., Horchler, B., Ha, N.-T., Reimer, C., Vieira, C.P., Vieira, J., Posnien, N., 2020. Multiple loci linked to inversions are associated with eye size variation in species of the Drosophila virilis phylad. Sci. Rep. 10, 12832. 10.1038/s41598-020-69719-z

Ridgway, A.M., Hood, E.J., Jimenez, J.F., Nunes, M.D.S., McGregor, A.P., 2024. Sox21b underlies the rapid diversification of a novel male genital structure between Drosophila species. Curr. Biol. CB 34, 1114–1121.e7. 10.1016/j.cub.2024.01.022

Ross-Innes, C.S., Stark, R., Teschendorff, A.E., Holmes, K.A., Ali, H.R., Dunning, M.J., Brown, G.D., Gojis, O., Ellis, I.O., Green, A.R., Ali, S., Chin, S.-F., Palmieri, C., Caldas, C., Carroll, J.S., 2012. Differential oestrogen receptor binding is associated with clinical outcome in breast cancer. Nature 481, 389–393. 10.1038/nature10730

Santos, M.E., Le Bouquin, A., Crumière, A.J.J., Khila, A., 2017. Taxon-restricted genes at the origin of a novel trait allowing access to a new environment. Science.

Sen, S., Reichert, H., VijayRaghavan, K., 2013. Conserved roles of ems/Emx and otd/Otx genes in olfactory and visual system development in Drosophila and mouse. Open Biol. 3, 120177. 10.1098/rsob.120177

Shannon, P., Markiel, A., Ozier, O., Baliga, N.S., Wang, J.T., Ramage, D., Amin, N., Schwikowski, B., Ideker, T., 2003. Cytoscape: a software environment for integrated models of biomolecular interaction networks. Genome Res. 13, 2498–2504. 10.1101/gr.1239303

Stampanoni, M., Groso, A., Isenegger, A., Mikuljan, G., Chen, Q., Bertrand, A., Henein, S., Betemps, R., Frommherz, U., Böhler, P., Meister, D., Lange, M., Abela, R., 2006. Trends in synchrotron-based tomographic imaging: the SLS experience, in: Bonse, U. (Ed.),. Presented at the SPIE Optics + Photonics, San Diego, California, USA, p. 63180M. 10.1117/12.679497

Streinzer, M., Brockmann, A., Nagaraja, N., Spaethe, J., 2013. Sex and caste-specific variation in compound eye morphology of five honeybee species. PloS One 8, e57702. 10.1371/journal.pone.0057702

Szklarczyk, D., Gable, A.L., Nastou, K.C., Lyon, D., Kirsch, R., Pyysalo, S., Doncheva, N.T., Legeay, M., Fang, T., Bork, P., Jensen, L.J., von Mering, C., 2021. The STRING database in 2021: customizable protein-protein networks, and functional characterization of user-uploaded gene/measurement sets. Nucleic Acids Res. 49, D605–D612. 10.1093/nar/gkaa1074

Tahayato, A., Sonneville, R., Pichaud, F., Wernet, M.F., Papatsenko, D., Beaufils, P., Cook, T., Desplan, C., 2003. Otd/Crx, a dual regulator for the specification of ommatidia subtypes in the Drosophila retina. Dev. Cell 5, 391–402. 10.1016/s1534-5807(03)00239-9

Torres-Oliva, M., Almudi, I., McGregor, A.P., Posnien, N., 2016. A robust (re-)annotation approach to generate unbiased mapping references for RNA-seq-based analyses of differential expression across closely related species. BMC Genomics 17, 392. 10.1186/s12864-016-2646-x

Torres-Oliva, M., Schneider, J., Wiegleb, G., Kaufholz, F., Posnien, N., 2018. Dynamic genome wide expression profiling of Drosophila head development reveals a novel role of Hunchback in retinal glia cell development and blood-brain barrier integrity. PLoS Genet. 14, e1007180. 10.1371/journal.pgen.1007180

Vandendries, E.R., Johnson, D., Reinke, R., 1996. orthodenticle is required for photoreceptor cell development in the Drosophila eye. Dev. Biol. 173, 243–255. 10.1006/dbio.1996.0020

Wakakuwa, M., Stavenga, D.G., Arikawa, K., 2007. Spectral organization of ommatidia in flower-visiting insects. Photochem. Photobiol. 83, 27–34. 10.1562/2006-03-03-IR-831

Wang, L.-H., Huang, Y.-T., Tsai, Y.-C., Sun, Y.H., 2010. The role of eyg Pax gene in the development of the head vertex in Drosophila. Dev. Biol. 337, 246–258. 10.1016/j.ydbio.2009.10.038

Warrant, E., Nilsson, D.-E. (Eds.), 2006. Invertebrate vision. Cambridge University Press, Cambridge, UKL; New York.

Warrant, E.J., 1999. Seeing better at night: life style, eye design and the optimum strategy of spatial and temporal summation. Vision Res. 39, 1611–1630. 10.1016/s0042-6989(98)00262-4

Weinkove, D., Neufeld, T.P., Twardzik, T., Waterfield, M.D., Leevers, S.J., 1999. Regulation of imaginal disc cell size, cell number and organ size by Drosophila class I(A) phosphoinositide 3-kinase and its adaptor. Curr. Biol. CB 9, 1019–1029. 10.1016/s0960-9822(99)80450-3

Xia, B., Zhang, W., Zhao, G., Zhang, X., Bai, J., Brosh, R., Wudzinska, A., Huang, E., Ashe, H., Ellis, G., Pour, M., Zhao, Y., Coelho, C., Zhu, Y., Miller, A., Dasen, J.S., Maurano, M.T., Kim, S.Y., Boeke, J.D., Yanai, I., 2024. On the genetic basis of tail-loss evolution in humans and apes. Nature 626, 1042–1048. 10.1038/s41586-024-07095-8

Zhang, Y., Liu, T., Meyer, C.A., Eeckhoute, J., Johnson, D.S., Bernstein, B.E., Nusbaum, C., Myers, R.M., Brown, M., Li, W., Liu, X.S., 2008. Model-based Analysis of ChIP-Seq (MACS). Genome Biol. 9, R137. 10.1186/gb-2008-9-9-r137

Zhou, Y., Zhou, B., Pache, L., Chang, M., Khodabakhshi, A.H., Tanaseichuk, O., Benner, C., Chanda, S.K., 2019. Metascape provides a biologist-oriented resource for the analysis of systems-level datasets. Nat. Commun. 10, 1523. 10.1038/s41467-019-09234-6

